# Action Potential Threshold Variability for Different Electrostimulation Models and the Impact on Occupational Exposure Limit Values

**DOI:** 10.1101/2024.04.22.590543

**Authors:** Florian Soyka, Thomas Tarnaud, Carsten Alteköster, Ruben Schoeters, Tom Plovie, Wout Joseph, Emmeric Tanghe

## Abstract

Occupational exposure limit values (ELVs) for body internal electric fields can be derived from thresholds for action potential generation. These thresholds can be calculated with electrostimulation models. The spatially extended nonlinear node model (SENN) is often used to determine such thresholds. An important part of these models are the membrane channel dynamics describing the ionic transmembrane currents. This work shows how ELVs change significantly with different ion channel dynamics (up to a factor of 22). Furthermore, two more detailed double-cable models by Gaines et al. (MRG-Sensory and MRG-Motor) are also considered in this work. Thresholds calculated with the SENN model (with Frankenhaeuser-Huxley membrane channel dynamics) and the MRG models are compared for frequencies between 1 Hz and 100 kHz and temperatures between 22 °C and 37 °C. Results show that MRG thresholds are lower than SENN thresholds. In the context of occupational ELVs, using the double cable model would lead to approximately ten times lower limit values. Therefore, future exposure guidelines should take the influence of different electrostimulation models into account when deriving ELVs.

**Highlights:**

1. Different membrane channel dynamics change derived exposure limit values by more than one order of magnitude.
2. Double-cable models result in a reduction of derived exposure limit values by one order of magnitude.
3. Lower temperatures reduce the action potential thresholds at frequencies below 300 Hz.

## INTRODUCTION

Low frequency magnetic fields can induce electric fields in human bodies. In turn, these electric fields can lead to the generation of action potentials in the nervous system. This type of electrostimulation is intended in some medical applications like TMS (transcranial magnetic stimulation). Besides that, however, it is considered an adverse health effect. Limit values have been introduced to restrict exposure of the general public and workers accordingly. Limit values exist for the body-internal electric fields as well as for the magnetic fields originating from some body-external source.

The International Commission on Non-Ionizing Radiation Protection (ICNIRP) is the main institution providing guidelines for limit values in Europe. They named the limits for the body-internal electric fields “basic restrictions” and for the magnetic fields “reference levels”. Nomenclature differs between standards. The Institute of Electrical and Electronics Engineers (IEEE) provides similar limits and named them “dosimetric reference limits” for the body-internal electric fields and “exposure reference levels” for the magnetic fields. The IEEE standard C95.1-2019 (IEEE Standards Coordinating Committee 39, 2019) is mainly used in America and Asia. Since the author’s institutions are based in Europe and partially deal with occupational exposure, this work will focus on the limits and the nomenclature given in the EU directive 2013/35/EU (The European Parliament and the Council of the European Union, 2013). However, similar reasoning applies for the limit values given by ICNIRP and IEEE for both the general public and occupational exposure.

The EU directive 2013/35/EU specifies occupational exposure limit values (ELVs) for adverse health effects for body-internal electric fields. These ELVs are based on recommendations given by ICNIRP for occupational exposure (International Commission on Non-Ionizing Radiation Protection, 2010). ICNIRP states that their recommendations are based on experimental findings as well as theoretical calculations using an axon model. The spatially extended nonlinear node (SENN) axon model (Reilly, et al., 1985) has been used by Reilly and Diamant (Reilly & Diamant, 2011) to derive exposure guidelines which are partially comparable to the ICNIRP guidelines and are being adopted by the IEEE standard C95.1-2019. Therefore, SENN model action potential thresholds play an important role in deriving ELVs.

Both ICNIRP and IEEE mention that more research is needed to improve the modelling accuracy (International Commission on Non-Ionizing Radiation Protection, 2020; Reilly & Hirata, 2016). Certain assumptions about SENN model parameters must be made to derive worst-case results which in turn guarantee maximum safety for the occupational setting. Deriving such assumptions is challenging and potential problems with the currently used parameters have been identified previously (Neufeld, et al., 2016). An important choice in the SENN model is the type of membrane channel dynamics (MCDs) that is being used. All the cited work so far is based on “Frankenhaeuser-Huxley” (FH) dynamics to model ionic membrane channel currents. Tarnaud et al. (Tarnaud, et al., 2018) presented SENN model results for four additional types of MCDs (Table 1). They showed that the results for pulsed waveforms can significantly differ between MCDs. In turn, this would lead to varying ELVs depending on which MCDs are being used. Therefore, this paper investigates how different MCDs influence action potential thresholds and in succession ELVs.

**Table 1:** The five models for membrane channel dynamics as described in Tarnaud et al. (2018). The FH model has traditionally been used for SENN calculations in the context of exposure guidelines. Data for the models were derived from different organisms.

| Model | Abbreviation | Experiments |
| --- | --- | --- |
| Frankenhaeuser-Huxley | FH | Frog node |
| Hodgkin-Huxley | HH | Squid axon |
| Chiu-Ritchie-Rogart-Stagg-Sweeney | CRRSS | Rabbit node |
| Schwarz-Eikhof | SE | Rat node |
| Schwarz-Reid-Bostock | SRB | Human nerve |

Another, potentially more realistic electrostimulation model than the SENN model was introduced by McIntyre et al. (McIntyre, et al., 2002) and is called the MRG model. The MRG model differs from the original SENN model by including paranodal sections, a double-cable structure, and altered membrane channel dynamics. For example, the MRG model was successfully used to describe experimentally determined perceptual thresholds for human arms and legs exposed to magnetic fields (Davids, et al., 2017). The MRG model was further refined by Gaines et al. (Gaines, et al., 2018) resulting in two models: “MRG - Sensory” and “MRG - Motor” which are more specific to the type of peripheral nerve under investigation.

These two models and the SENN model with FH dynamics are implemented in the Sim4Life^1^ simulation environment and were used to determine action potential thresholds for this study. Furthermore, the freely available SENN model implementation by Reilly & Diamant (Reilly & Diamant, 2011) and another freely available SENN model implementation (called EONS^2^) by Tarnaud et al. (Tarnaud, et al., 2022) were used. Therefore, there were five simulation setups in total: (i) SENN by Reilly & Diamant, (ii) SENN in EONS, (iii) SENN in Sim4Life, and (iv) MRG - Sensory and (v) MRG - Motor in Sim4Life as well.

The goal and novelty of this study was to compare action potential thresholds and the potential ELVs resulting from these thresholds for different electrostimulation models. Furthermore, the influence of temperature on action potential thresholds was investigated.

The methods section starts with verification of our SENN model implementation (EONS) for the FH dynamics by comparing action potential thresholds to the values originally found by Reilly & Diamant. Furthermore, it is shown how ELVs can be derived from action potential thresholds and which safety factors must be accounted for. Next, the models and their setup are described. The results section is divided into two parts. First, ELVs derived for five different MCDs are presented. Second, differences between SENN and MRG model results are shown, as well as the temperature dependency of the two models. Subsequently, the discussion compares the results of this study to previous work and highlights the potential impact of the findings on future safety guidelines.

Note, that this article is based on two previous extended abstracts presented at BioEM conferences (Soyka, et al., 2022; Soyka, et al., 2023).

## METHODS

To investigate the different MCDs, the SENN model implementation in MATLAB (EONS) from Tarnaud et al. (Tarnaud, et al., 2018) was used. The model parameters were adjusted to match the original parameters used for deriving exposure guidelines by Reilly & Diamant, while also including finite myelin impedance (number of myelin layers, conductance and capacitance per layer are adopted from (Gomez-Tames, et al., 2019)). Note that in this case both models use the FH MCDs. Figure 1 shows that apart from small discrepancies for low frequencies the model results match very well. These small discrepancies can be attributed to differences in implementation (e.g., used discretization scheme and tolerances) and the explicit presence of internodes with finite myelin impedance in EONS. Therefore, the EONS model provides comparable results and can be used for further calculations.

**Figure 1:**
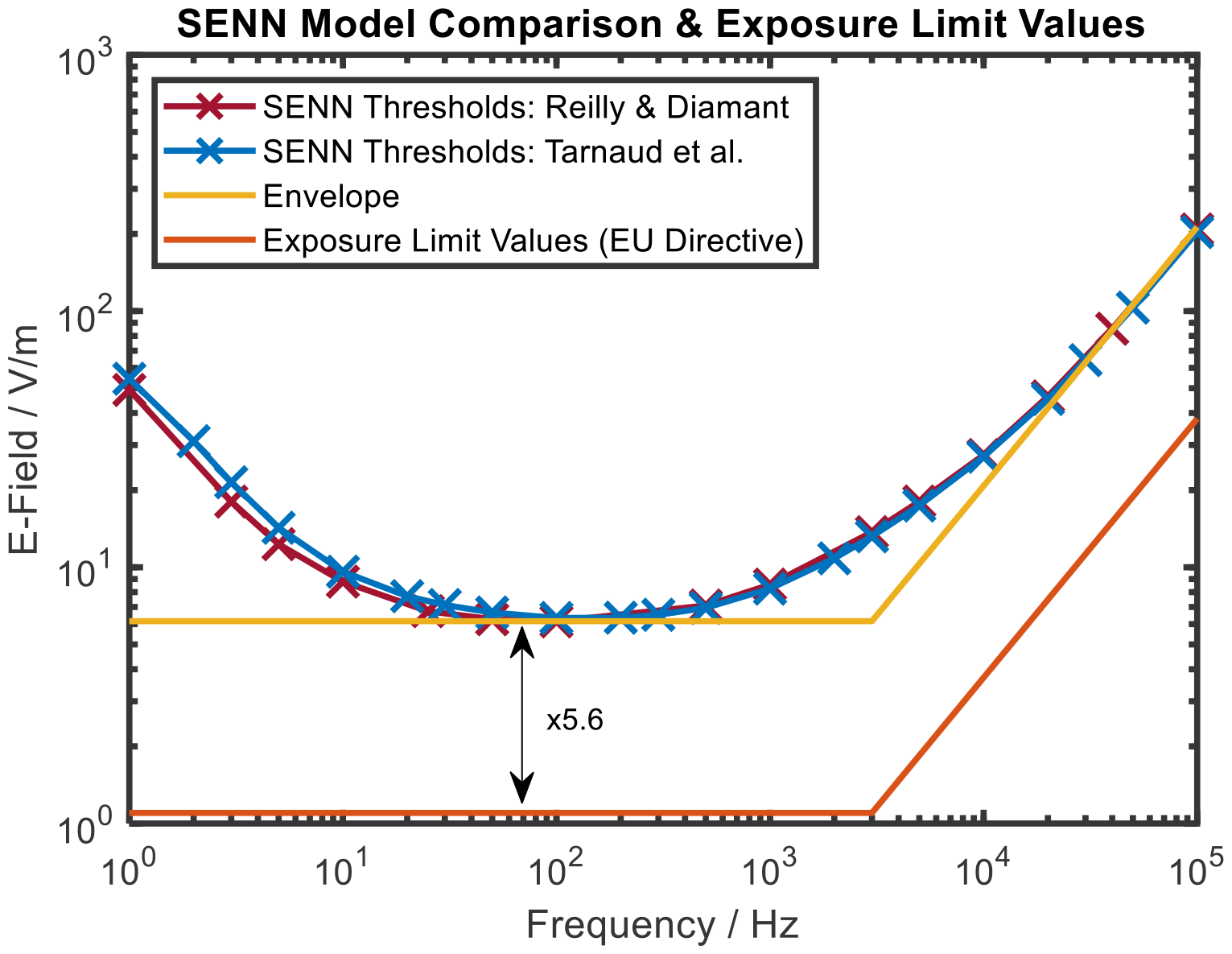
SENN model thresholds for action potential generation (red & blue) are given for sinusoidal waveforms as a function of frequency for two different SENN model implementations (Reilly & Diamant and the EONS model by Tarnaud et al.). The results are in good agreement. An envelope can be found from which the exposure limit values can be derived by applying a safety factor of 5.6. The temperature was 22 °C.

For a spatially uniform time-varying electric field parallel to a straight finite myelinated axon with sealed terminals, the SENN model calculates the minimum electric-field amplitude at which an action potential is elicited. Figure 1 shows the excitation thresholds for sinusoidal fields of different frequencies. The ELVs can be derived from the SENN thresholds by placing an envelope around the thresholds and applying a safety factor. The envelope consists of two lines. The first line has a constant value E_0_ in V/m which is defined by the lowest threshold. The second line is proportional to the frequency (i.e., slope equal to unity in the logarithmic plot, in agreement with the international guidelines/standards (The European Parliament and the Council of the European Union, 2013; IEEE Standards Coordinating Committee 39, 2019; International Commission on Non-Ionizing Radiation Protection, 2010), and with the strength-frequency curve of the SENN-model (Reilly, 2012, pp. 134-141)). It is starting from the corner frequency f_C_ which is chosen such that all thresholds are just enclosed by the envelope. The safety factor between the ELVs from the EU directive and the SENN threshold envelope is 6.15 V/m / 1.1 V/m ≈ 5.6. Note that the ELVs are defined up to a frequency of 10 MHz, but that the current study investigates thresholds only up to 100 kHz.

Using the EONS SENN model with the original parameters used for deriving exposure guidelines by Reilly & Diamant, thresholds were calculated for 5 MCDs (Hodgkin & Huxley, 1952; Frankenhaeuser & Huxley, 1964; Chiu, et al., 1979; Sweeney, et al., 1987; Schwarz & Eikhof, 1987; Schwarz, et al., 1995). The model labels and abbreviations are given in Table 1 and further information about them can be found in Tarnaud et al. (Tarnaud, et al., 2018).

The SENN model implementation by Reilly & Diamant represents the standard to which the other models can be compared, because its results form the basis for the current ELVs. Reilly & Diamant chose a temperature of 22 °C for their studies. This temperature cannot be adjusted in their software, without modifying and recompiling the FORTRAN source code which we did not do in this study. In the Sim4Life and EONS simulation tools temperature settings can easily be adjusted. The body core temperature of 37 °C was chosen as a comparison value to investigate the influence of temperature on the thresholds. The Sim4Life simulation environment allows calculating thresholds for the SENN FH and the two MRG models for both temperatures. However, in some cases no valid thresholds could be obtained with the Sim4Life models for very low or very high frequencies. Therefore, the EONS SENN FH model implementation was used in addition which allowed the calculation of SENN model thresholds for both temperatures across the full frequency range. Furthermore, having three different simulation tools allows for a cross comparison between the tools for the SENN model at 22 °C.

All simulations used the same setup (matching the original setup by Reilly and Diamant): an axon (20 µm diameter) within and parallel to a homogenous electric field with a sinusoidally varying amplitude (a sine with frequencies from 1 Hz to 100 kHz). The simulation tools adjust the amplitudes via a titration procedure until the smallest amplitude (the threshold) is found for which an action potential (at least 80 mV depolarization of three consecutive nodes) is elicited (Reilly & Diamant, 2011). After calculating all five simulation setups at 22 °C, all but Reilly & Diamant’s SENN model implementation were additionally run at 37 °C.

In accordance with a worst-case approach, the procedure described above for deriving ELVs was applied for the lowest thresholds found from the previous calculations. The resulting ELVs are compared to the current ELVs from the EU directive.

The time course of the membrane voltage at the first and last node of Ranvier was visually checked at threshold level intensity to verify the action potential. For some frequencies the time course followed the sinusoidal shape of the stimulus and did not show the typical action potential shape in the Sim4Life simulation environment. These cases were excluded from the results. Please note that the excluded cases are not essential for answering our research questions.

## RESULTS

### EXPOSURE LIMIT VALUES DERIVED FOR DIFFERENT MEMBRANE CHANNEL DYNAMICS WITH THE SENN MODEL

Figure 2 shows calculations of SENN model thresholds for five MCDs. The results for the FH dynamics are the same as in Figure 1 and can be used to derive the current ELVs given in the EU directive (The European Parliament and the Council of the European Union, 2013).

**Figure 2:**
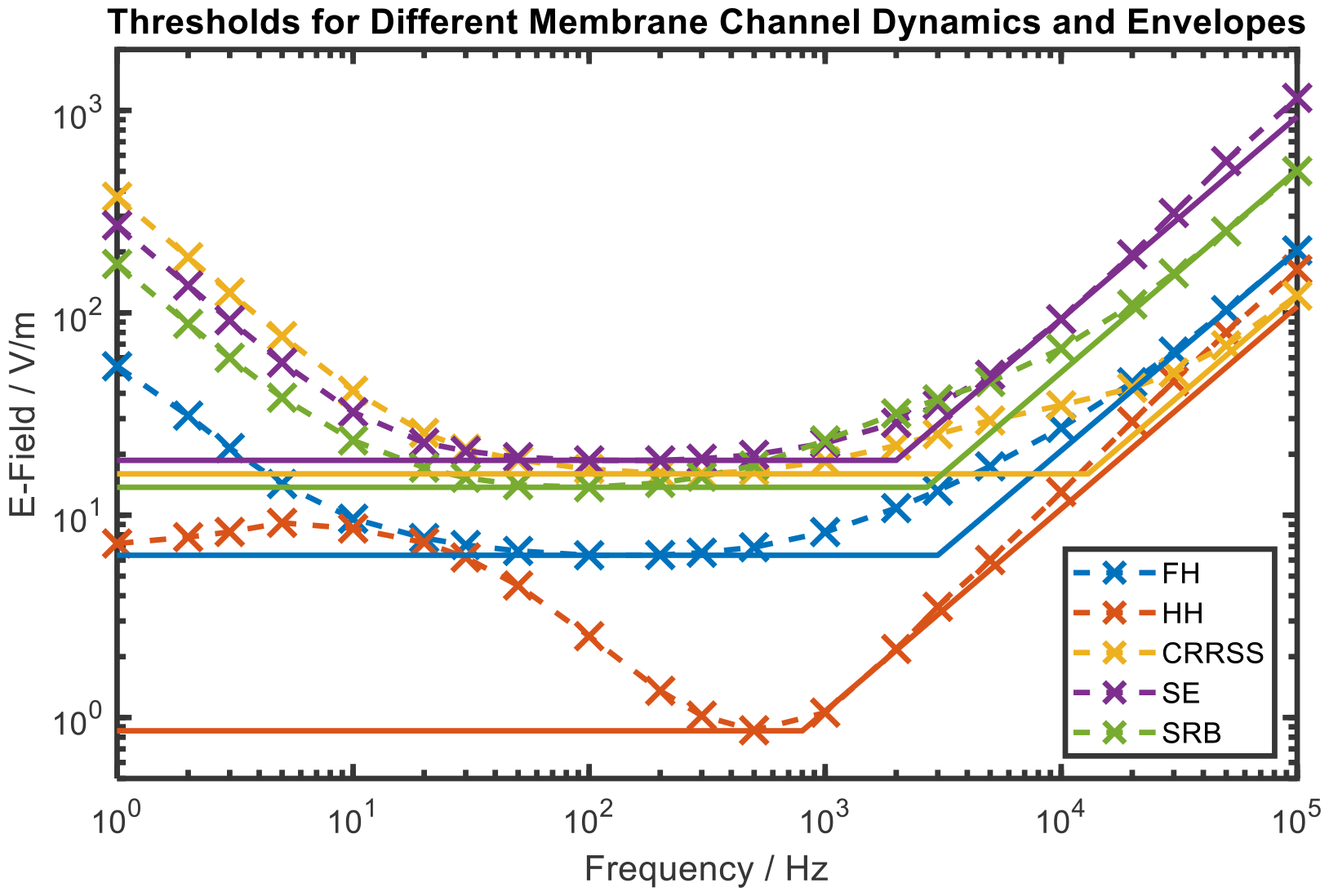
SENN model thresholds for five MCDs calculated with EONS (see Table 1 for abbreviations). The results for the FH dynamics correspond to Figure 1. Envelopes were derived for all dynamics and the resulting parameters can be found in Table 2. The temperature was 22 °C.

Envelopes for the remaining four MCDs were derived according to the description in the Methods. Table 2 lists the respective parameters E_0_ and the corner frequency f_C_. Exposure limit values can further be derived from these envelopes by applying a safety factor of 5.6 (Figure 3).

**Table 2:** Parameters for the five envelopes which were derived from the SENN model thresholds for the different MCDs at 22 °C.

| MCD Model | $E_0$ / V/m | Corner Frequency $f_c$ / kHz |
| --- | --- | --- |
| HH | 0.86 | 0.8 |
| FH | 6.34 | 3.0 |
| CRRSS | 16.0 | 13.0 |
| SE | 18.66 | 2.0 |
| SRB | 13.74 | 2.7 |

**Figure 3:**
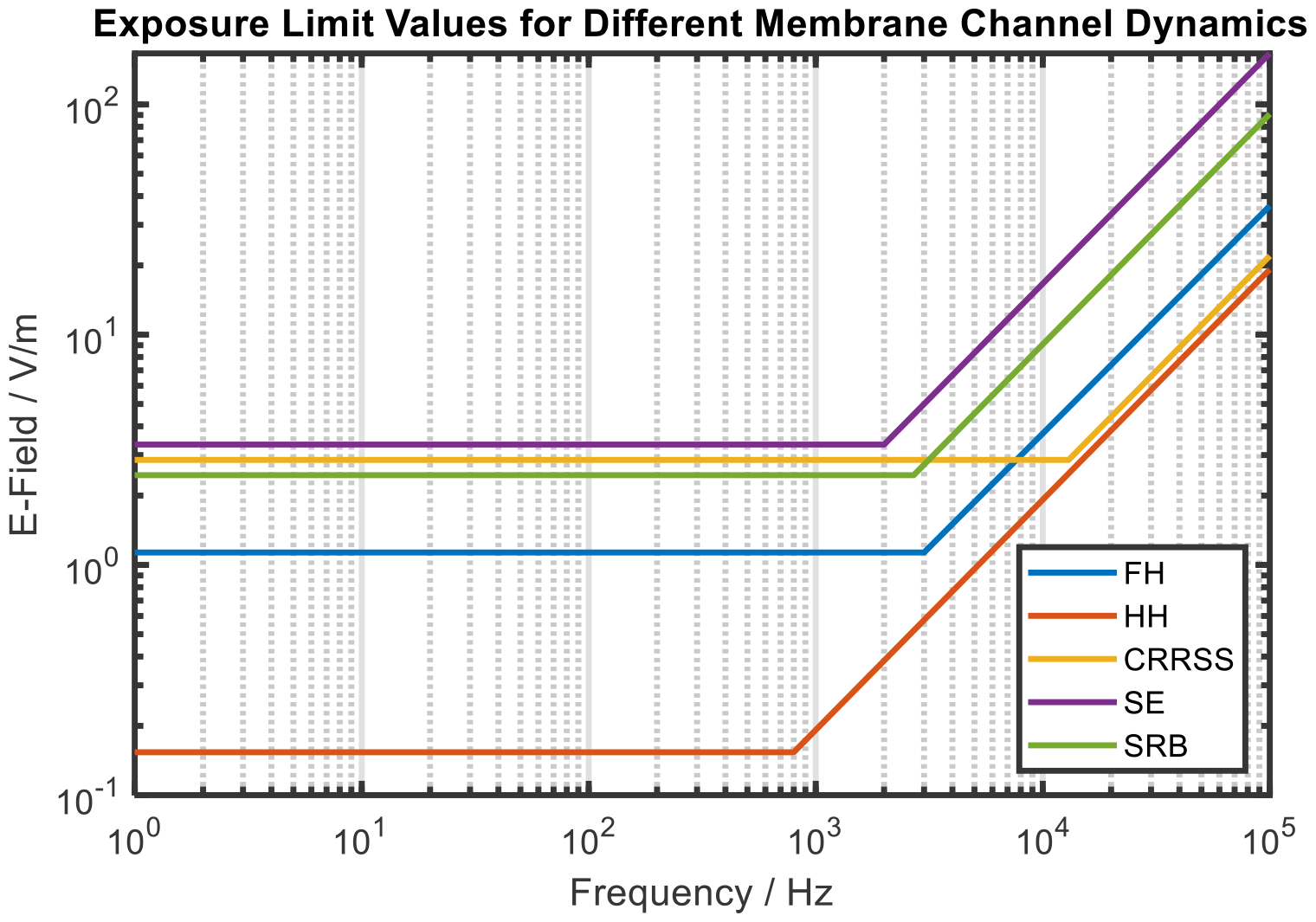
Exposure limit values derived from SENN model thresholds for different membrane channel dynamics (Figure 2). The values for the Frankenhaeuser-Huxley dynamics correspond to the current values given in the EU directive. ELVs can differ up to a factor of 22 between different MCDs. The temperature was 22 °C.

### COMPARISON BETWEEN SENN AND MRG MODEL RESULTS AND IMPACT OF TEMPERATURE

Figure 4 shows the thresholds for the five simulation setups at 22 °C. The SENN thresholds are very similar for all simulation tools and only show negligible differences in the low frequency range. The Sim4Life simulation environment had difficulties finding the thresholds for some frequencies and therefore these values were excluded (missing yellow markers). Figure 5 shows the same thresholds in addition with the calculations at 37 °C for all but Reilly & Diamant’s model. The thresholds are significantly increased below 300 Hz for all models (largest increase with a factor of 7.3 for EONS SENN), while for frequencies above 300 Hz the impact of temperature is smaller (<30%).

**Figure 4:**
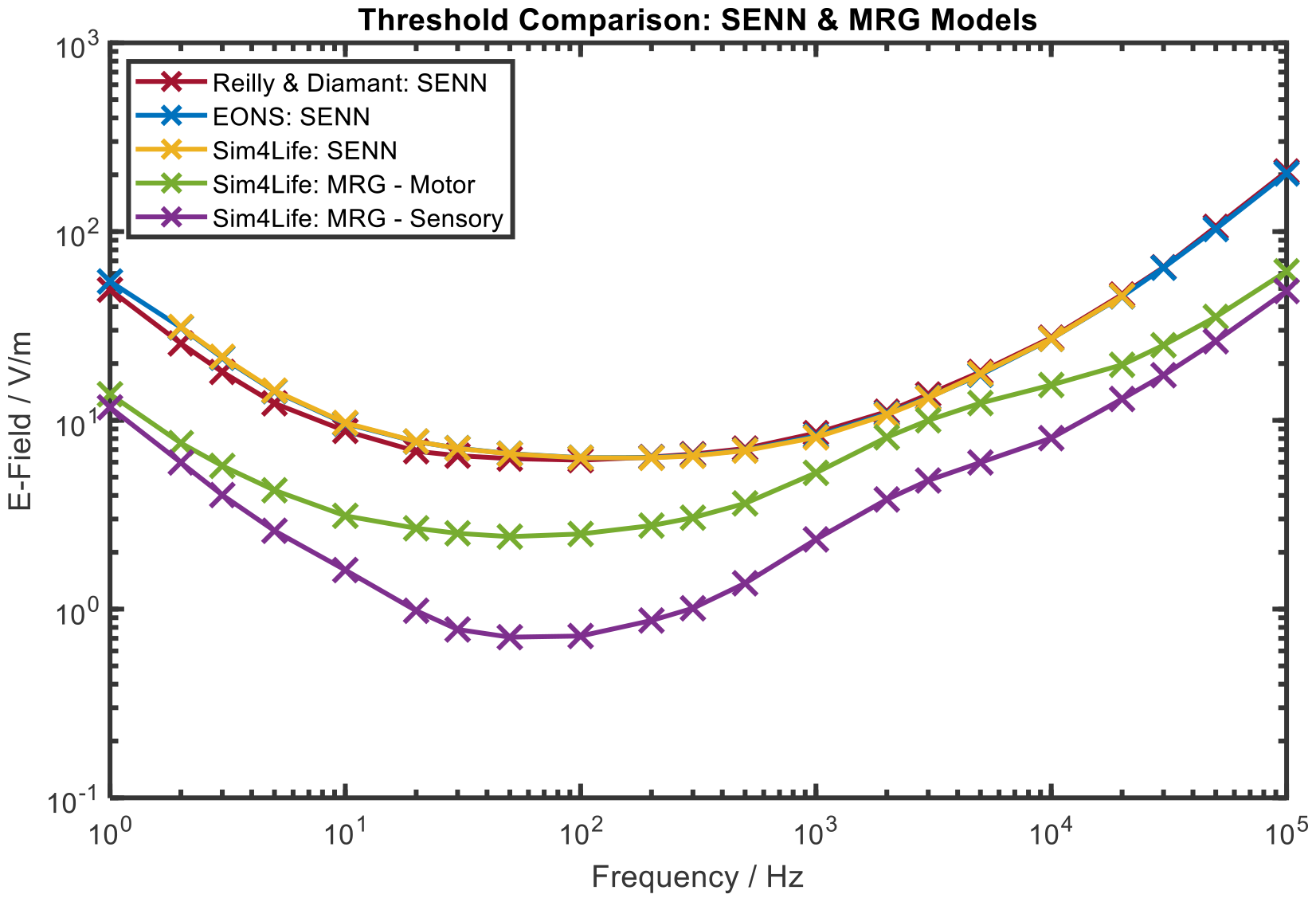
Thresholds for action potential generation based on five simulation setups for a temperature of 22 °C. The SENN model thresholds are very similar for the three different simulation tools. The MRG – Sensory model produces the lowest thresholds.

**Figure 5:**
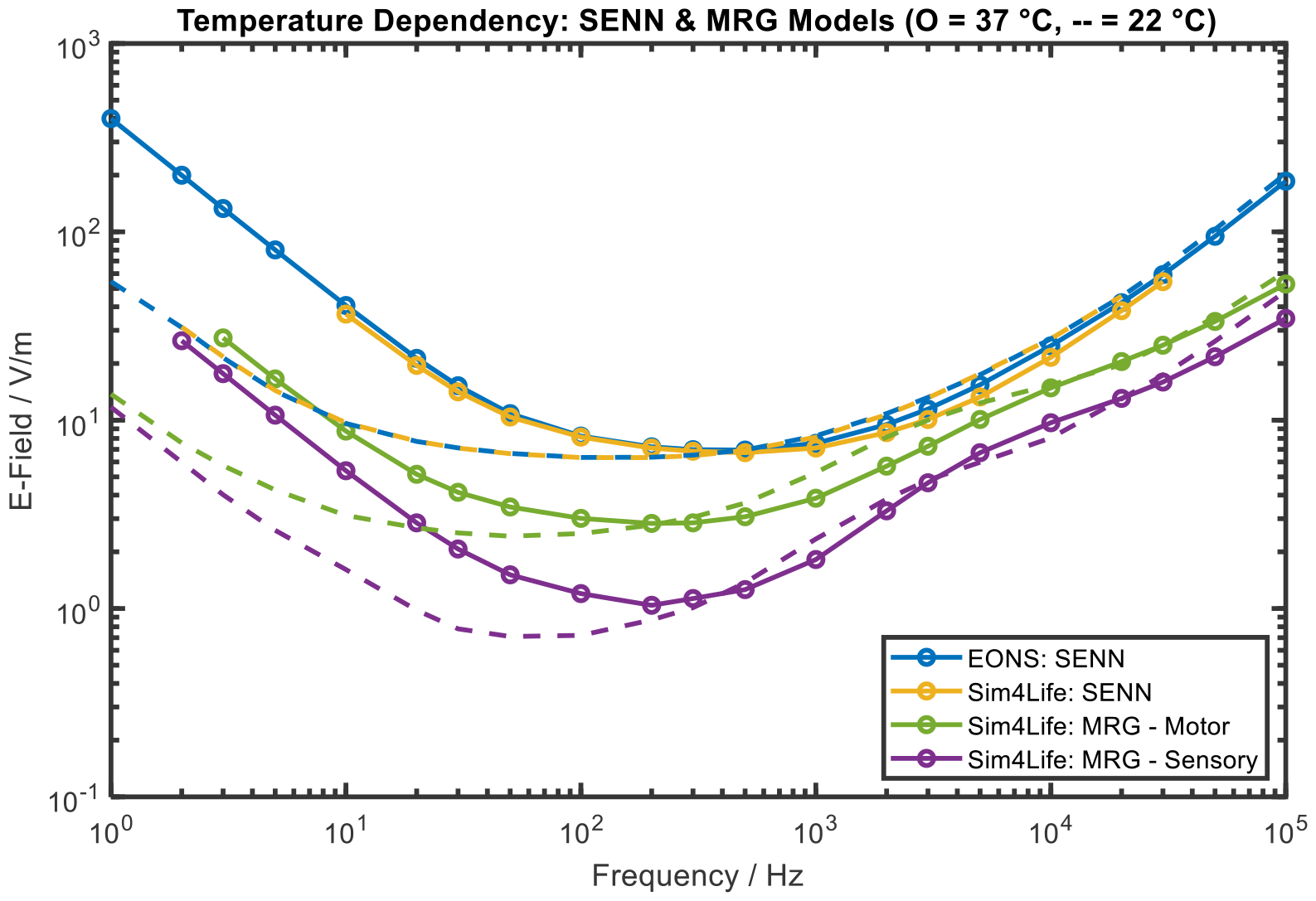
Thresholds are shown for two temperatures: 22 °C (dashed lines, same as in Figure 4) and 37 °C (solid line and circular markers). The EONS and the Sim4Life simulation tools give very similar results for the SENN model. For frequencies below 300 Hz temperature has a significant influence, resulting in higher thresholds for higher temperatures.

Figure 6 shows the thresholds for the EONS SENN model and the MRG - Sensory model for both temperatures, together with the corresponding envelopes and the resulting ELVs. The MRG - Sensory model was chosen because it has the lowest thresholds and can be seen as a worst-case scenario from an occupational safety perspective. The EONS SENN model is comparable to Reilly & Diamant’s original model (Figure 1) and additionally allows investigating the influence of temperature on thresholds. It represents the currently implemented ELVs in the EU directive.

**Figure 6:**
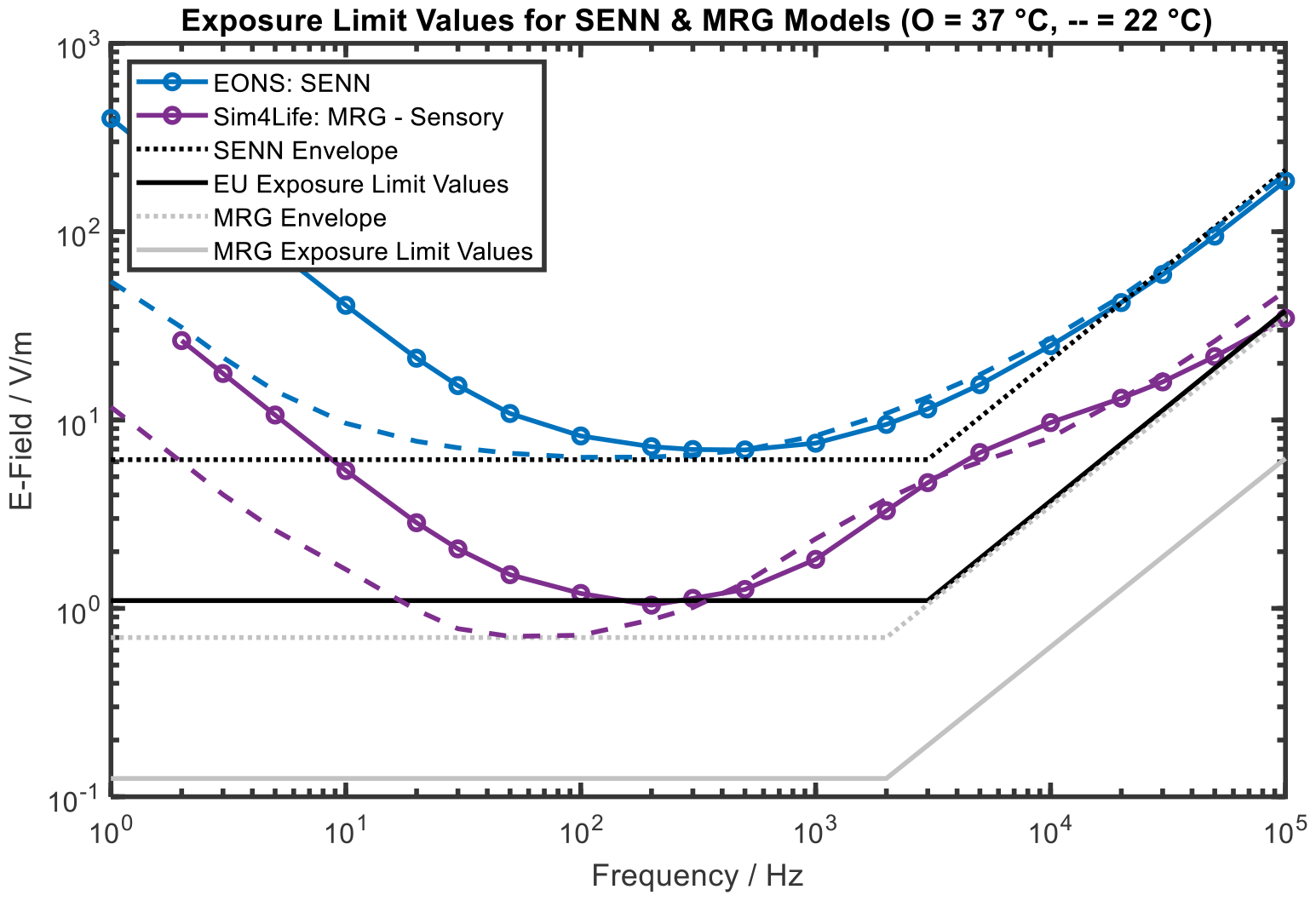
Thresholds for the SENN (FH) and the MRG – Sensory models at 22 °C and 37 °C. The corresponding envelopes (dotted lines) are used to derive the ELVs which include an additional safety factor of 5.6. The ELVs based on the MRG model would be up to 10 times lower than the ELVs given in the current EU directive. Note that thresholds at 22 °C provide the most conservative values for defining the envelopes.

The envelope for the EONS SENN model has the following parameters: lowest threshold *E*_0_ = 6.3 V/m and corner frequency *f*_*C*_ = 3 kHz. And the envelope for the MRG - Sensory model has the following parameters: lowest threshold *E*_0_ = 0.7 V/m and corner frequency *f*_*C*_ = 2 kHz.

## DISCUSSION

Figure 3 shows that potential ELVs can differ significantly depending on the MCDs being used. The most pronounced difference can be found at 1 Hz between the HH and the SE dynamics and corresponds to a factor of 22. Note that the differences are smaller at high frequencies because the HH dynamics have a lower corner frequency. Since the current ELVs in the EU directive are based on the FH MCDs they are rather conservative. They are smaller than the SE and SRB values, as well as the low frequency CRRSS values. Due to the safety factor of 5.6 the FH based ELVs are lower than the high frequency thresholds for the HH and CRRSS dynamics (Figure 2). However below 3 kHz the HH thresholds are lower than the FH based ELVs. In summary, the current (FH based) ELVs might be overly conservative or not conservative enough, depending on which MCDs are used to derive them. This raises the question which MCDs are most realistic given experimental data (Reilly & Hirata, 2016; Soldati, et al., 2018; Suzuki, et al., 2022).

The five MCDs used in this study are the first published Hodgkin-Huxley type ion channel models, derived from voltage-clamp measurements (see Table 1). Consequently, these classical MCD models have been used in numerous neurostimulation models. For example, for the application of cochlear nerve modeling, the first four membrane models (HH, FH, CRRSS and SE) are compared in Cartee et al. (Cartee, et al., 2000) and O’Brien & Rubinstein (O’Brien & Rubinstein, 2016). These are all valid models and since the human nervous system includes many different types of neurons, a large variability of channel dynamics is present in humans as well. Experimental data for thresholds in humans is sparse and therefore a direct comparison is difficult. Our research is in line with a survey comparing ten numerical electrostimulation models (Reilly, 2016), which found significant differences between thresholds depending on the model. More research is needed, especially in the direction of experimental validation of the models and that other MCDs besides the FH dynamics should be considered as well. This work established that the choice of MCDs plays an important role for the derivation of the ELVs.

Looking at the behavior of thresholds as a function of frequency (Figure 2) some differences can be seen between the MCDs. For example, the HH thresholds are stable for low frequencies whereas the thresholds for all other MCDs decrease. Further calculations showed that these differences are not present in the strength-duration curves obtained with rectangular waveforms. More research is needed to better understand this behavior. This example also highlights the importance of investigating the effect of non-sinusoidal waveforms for different MCDs since the action potential generation can be significantly different. To facilitate the analysis of different MCDs and the effects of non-sinusoidal waveforms, we propose the EONS (Evaluation of Non-Sinusoidal Magnetic Fields) software in Tarnaud et al. (Tarnaud, et al., 2022).

Comparing the SENN model thresholds at 22 °C (Figure 4) shows that all three simulation tools give very similar results. The same holds true at 37 °C for the Sim4Life and the EONS simulations (Figure 5). This cross-check between simulation tools provides a good indicator for the validity of the simulations. One of the goals of this study was to investigate the influence of temperature on the action potential thresholds. Figure 4 shows that for frequencies above approximately 300 Hz the thresholds for 22 °C and 37 °C are rather similar. Below 300 Hz the thresholds for 37 °C are significantly higher than those for 22 °C (e.g., 7.3 times higher for EONS-SENN at 1 Hz). An increase of the thresholds with temperature is expected at low frequencies, because the ion channels’ time constants become smaller at higher temperatures. At low frequencies, the effect of fast sodium activation can be neglected, compared to the relatively slow inactivation and activation of sodium and potassium currents, respectively. As a result, sodium current inactivation and potassium current activation will increasingly counteract neuronal excitation at higher temperatures, eventually resulting in heat block (Rattay & Aberham, 1993; Mou, et al., 2012). In contrast, at high frequencies the sensitivity of the threshold to the temperature is small (<30% discrepancy between the 22 °C and 37 °C thresholds above 300 Hz), because all the gate parameters will need several periods to reach their steady state values. As thresholds at 22 °C are lower than at 37 °C, they provide a conservative estimate for potential adverse health effects. Note that current ELVs are based on SENN thresholds at 22 °C (Figure 6). For future guidelines, it might be helpful to consider the temperature differences between body parts e.g., the limbs in comparison to the chest or the brain.

In general, it was found that the MRG models give up to ten times lower thresholds and ELVs than the SENN model calculations, in agreement with other computational exposure studies (Reilly, 2016; Neufeld, et al., 2016). This is of course very important from a safety point of view since it raises the question if the current ELVs are too high. For example, Figure 6 shows that the MRG – Sensory thresholds around 50 Hz are below the current ELVs. The thresholds for the MRG models have inflection points around 3 kHz which are not present for the SENN model thresholds. Therefore, an envelope rising linearly with increasing frequency might not be the best option for deriving ELVs from these thresholds. Since both MRG models show these inflection points for both temperatures, it seems likely that this is a real effect and not some kind of artifact. Indeed, the sensory and motor MRG models include active membrane dynamics in the paranodal and internodal sections. Inflection points are expected when different frequencies result in initiation of action potentials at different locations along the axon, similar to observed deviations from classical strength-duration curves (Rattay, et al., 2012).

It is very important to get good experimental data to be able to verify and choose between different models. Davids et al. describe a good fit for perceptual thresholds in arms and legs for magnetic field stimulation between approximately 0.5 and 10 kHz with the original MRG model (Davids, et al., 2017). Fresnel et al. are planning on running similar studies at 50 Hz and 60 Hz for magnetic field stimulation of the leg (Fresnel, et al., 2022). Future work could pool such data and try to differentiate between different models. However, such an approach needs to model the induction of the electric field in the body as well. Properly calculating the induced electric field is important because the electrostimulation depends on the orientation of the nerve fibre with respect to the field.

## CONCLUSION

This work showed how the current ELVs given in the EU directive can be directly derived from SENN model action potential threshold calculations. Changing the membrane channel dynamics used in the SENN model leads to significantly different threshold values and in turn to different ELVs (up to a factor of 22). Depending on which MCDs are being used the current ELVs can be seen as either too conservative or as not conservative enough. Furthermore, this work showed that action potential thresholds based on MRG electrostimulation models are lower than thresholds calculated by the SENN model. Calculating thresholds for temperature settings of either 22 °C or 37 °C only made a difference for frequencies smaller than 300 Hz. Deriving exposure limit values from MRG thresholds in the same way as they are derived from SENN thresholds, results in approximately 10 times smaller values than those currently given in the EU directive for occupational exposure.

Future work will consist of gathering experimental data, which is needed to understand which model is better suited to derive exposure guidelines. Future guidelines should take the effect of different MCDs in the SENN model as well as the MRG model results into account.

## Data availability statement

The data that support the findings of this study are available from the corresponding author upon reasonable request.

The EONS simulation tool is freely available on GitHub: https://github.com/Florian-Soyka/EONS

## Footnotes

1 Sim4Life V7.0, ZMT, https://zmt.swiss/sim4life/

2 https://github.com/Florian-Soyka/EONS

